# Prediction Analysis of Preterm Neonates Mortality using Machine Learning Algorithms via Python Programming

**DOI:** 10.1101/2023.01.20.524905

**Authors:** Vahid Monfared, Afkham Hashemi

**Affiliations:** Postdoctoral Research Fellow, Division of Engineering in Medicine, Brigham and Women’s Hospital, Harvard Medical School, Cambridge, MA, USA; Department of Mechanical Engineering, Zanjan Branch, Islamic Azad University, Zanjan, Iran; Department of Electrical and Computer Engineering, Semnan University, Semnan, Iran

**Keywords:** Survival prediction, preterm, neonate, mortality, machine learning

## Abstract

Prediction analysis of preterm neonate mortality is necessary and significant for benchmarking and evaluating healthcare services in Hospitals and other medical centers. Application of artificial intelligence and machine learning models, which is a hot topic in medicine/healthcare and engineering, may improve physicians’ skill to predict the preterm neonatal deaths. The main purpose of this research article is to introduce a preterm neonatal mortality risk prediction by means of machine learning/ML predictive models to survive infants using supervised ML models if possible. Moreover, this paper presents some effective parameters and features which affect to survive the infants directly. It means, the obtained model has an accuracy of about 91.5% to predict the status of infant after delivery. After recognizing the critical status for an infant, physicians and other healthcare personnel can help to infant for possible surviving using special medical NICU cares. It has been tried to get some suitable models with high accuracy and comparing the results. In a word, a survival prediction analysis of preterm neonate mortality has been carried out using machine learning methods via Python programming (possible surviving infants after delivery in the hospital).

## 1. Introduction

Here, some highlighted research works have been introduced regarding our study. The neonatal period (1^st^ 28 days of life) is the phase of developing physiological adaptations for extra-uterine life. Preterm births (PTB) may be resulted in the important long-term neurodevelopmental difficulties and issues, pulmonary dysfunction, and visual impairment [1,2].

The aim of article [3] is to assess the result for wholly infants born before 33 weeks gestation till discharge from hospital. Probabilities of survival varies importantly in accordance with length of gestation among very preterm babies. A huge and considerable proportion of the deaths are related to a decision to limit intensive care at all ages of pregnancy [3]. One of the important purposes of a study [4] was to obtain survival to discharge for a great Canadian cohort of preterm infants admitted to the NICU (neonatal intensive care unit), and also to survey the effect of gender on survival and the effect of increasing postnatal age on predicted survival. Actuarial analysis provides beneficial information once counseling parents and highlights the importance of often revising the prediction for long-term survival specially after the first few days of life [4]. PTB (Preterm births) has been enlarged around 30% since 1981 and included 12.5% of wholly births in year 2004 in United States of America [5].

In industrial countries, preterm delivery is answerable for 70% of neonatal mortalities and 75% of morbidities. For example, in some countries like IRAN, the most common reason of neonatal death is also prematurity which is considered for 57.07% of neonatal death [6]. Low birth weight (LBW) is also accounted for two third of neonatal deaths [7]. As well as some infants born before 37 weeks of gestation are at bigger risk for morbidity and mortality in comparison with term newborns [8]. The considerable development in survival in France for newborns born at 25 out of 31 weeks’ gestation was accompanied by a significant decreasing in severe morbidity, but survival remained rare before 25 weeks. Though improving in survival at enormously low gestational age can be probable, its effect on long-term outcomes needs additional investigations. The long-term outcomes of the EPIPAGE-2 study will be instructive about this [9].

In the different study, the authors [10] estimated the frequency of maternal and perinatal results in women with diverse categories of preterm and term births, factors associated with lesser perinatal outcomes and related management interventions. Make a decision based on prognostic models may result to the care given to the premature infant being more individualized and with a improved use of resources. Predictive algorithms and models of mortality in premature neonates in Spain require to be extended [11].

The study [12] presented numerous risk factors for death among preterm neonates. A large number of the risk factors are generally avoidable. So, it is vital to address neonatal and maternal factors identified in the study [12] through suitable ANC and optimal infant medical care and feeding practices for reduction the high rate of preterm death. To illustrate survival and main morbidity among infants born tremendously preterm in China over the past decade. The researchers [13] found that infants born enormously preterm were at increased risk of mortality and morbidity in China, with a survival rate that improved over time and a main morbidity rate that increased. These findings propose that more active and effective treatment strategies are required, particularly for infants born at gestational age 25 to 27 weeks [13].

The results [14] proved that the DeepPBSMonitor model outperforms other methods, with an accuracy, recall, and AUC score of 0.88, 0.78, and 0.89, respectively. The suggested model has confirmed efficacy in predicting the real-time mortality risk of preterm infants in initial NICU hospitalization. The decision tree and Bayesian ridge imputation are implemented utilizing the SKLEAREN package in Python open-source software. For many imputations, the parameter sample posterior is set to be true and different random states are applied for each imputed dataset. Monochorionicity and GWG <10 kg are two main risk factors for PTB (Preterm births) before 32, 34, and 37 weeks, while maternal age, PE, and ICP were also risk factors for PTB in specific gestational age [15]. It should be mentioned that the firstseven7 days of admission is the danger time to death with median time of six days. Being born to antepartum hemorrhage mother, lack of Kangaroo mother care, unable to start feeding with 24-hr, Apnea and dehydration are the effective features and predictors of time to death. Thus, intervention that focuses on the identified predictors could have a paramount effect to prolong time to death and decrease preterm mortality [16].

This research has been done and conducted in Tehran, Iran in two phases. Firstly, critical risk factors in neonatal death were identified and then numerous machine learning models including RF (random forest), SVM (support vector machine), and LR (logistic regression), XGBClassifier, and Ensemble models were developed. Finally, these models were applied to predict neonatal death in a NICU and followed up the neonates to compare the results and outcomes of these neonates with real outcomes logically.

In this study, it has been tried to analyze and predict the final status (dead or alive) of an infant immediately after delivery in hospital, as well as finding important features and investigating (discussing) the accuracy of the AI/ML models. Neonatal results within dissimilar gestational ages and risk factors for neonatal mortality are analyzed in numerous countries, however a wide-ranging dataset is not accessible in IRAN, and so it is hard for healthcare professionals to provide reliable information for parents regarding outcomes of the preterm infants. Thus, with having a suitable knowledge regarding the result, influences the professionals to take care of mothers and their infants. In a word, the major aim is to predict and evaluate neonatal mortality rate of preterm infants along with analysis of the predictors (features) and risk factors using ML/AI in Iranian population which may be extended and beneficial to the other countries.

## 2. Material and Method

This study had no financial support. It comprised all preterm (26-37 weeks) infants (samples ∼ 1610) born alive in Hospital, throughout one-year period. The mentioned datasets were gathering utilizing questionnaires from patient medical records. The survey includes of fetal-neonatal, maternal, and pregnancy data; antenatal medications; root of delivery; management in the neonatal period; postnatal outcomes (described below) and delivery to discharge duration. Survey has been done for each preterm baby throughout the study period, (N∼1610). It means that our dataset has a shape in the form of (1610,30). Inclusion criteria was all live born preterm (before 37 weeks of gestation) babies delivered during the study period, as a minimum one examination in first 20 weeks (for precise estimation of GA). Infants born from diabetic and hypertensive mothers, who had lethal congenital anomalies, who did not receive prenatal steroid (one or two doses of 12 mg Betamethasone, 24 hours apart) at the time of admission and who received antenatal magnesium sulfate were excluded from the analysis.

Main consequence was delivery to discharge survival rate among preterm infants. Initial prognostic variables for survival are including, ‘Number_of_Fetus’, ‘Presentation’ (leading anatomical part of the fetus), ‘Mathernal_Level_of_Education’, ‘Mathernal_Age_Year’, ‘Gravidity’ (number of pregnancies), ‘Parity’ (number of deliveries), ‘History_of_Abortion’ (pregnancy terminated under 20 weeks), ‘Number_of_Live_Children’, ‘History_of_Child_Death’, ‘Delivery_Method’, ‘Indication_of_Cesarean_Delivery’, ‘Pregnancy_Complication’, ‘Indication _of_Termination’, ‘Fetal_Gender’, ‘Need_for_CPR’, ‘Need_for_NICU_Admition’ (defined as admitting in NICU for more than Twelve hours), ‘Corticosteroid_Administration’, ‘Surfactant_ Administration’ (administered in cases of respiratory distress syndrome), ‘Fetal_Anomaly’ (including cardiovascular, urogenital, skeletal, gastrointestinal and central nervous system anomalies), ‘STATUS_dead_alive’, ‘First_Minute_Apgar’, ‘Fifth_Minute_Apgar’, ‘Gestational_ Age_LessEqual_28weeks’, ‘Gestational_Age_LessEqual_32weeks’, ‘Gestational_Age_Less Equal_34weeks’, ‘Birth_Wieght_gr’, ‘Birth_Weight_LessEqual_2500gr’, ‘Birth_Weight_Less Equal2000gr’, ‘Birth_Weight_LessEqual_1500gr’, ‘Birth_Weight_LessEqual_1000gr’, briefly (30 variables). It should be mentioned that APGAR score is calculated by neonate’s color, heart rate, reflex irritability, muscle tone and respiration. It means, we have 29 inputs (features and attributes) and one output (label: STATUS_dead_alive). The main purpose is first learning and then predicting of new datasets using the best predictive model (dataset shape: (1610, 30)).

GA (Gestational_Age) was determined by last menstrual period and sonography in the first half of pregnancy. Apgar score was obtained on the basis of mother’s sex and GA was achieved by the neonatal birth weight and gestational diabetes. According to current guidelines developed by the American Academy of Pediatrics, we did not initiate resuscitation for infants younger than 23 weeks or those whose birth weight was less than 400 gr [17]. Survival was defined as the proportion of live births surviving up to the time of discharge. Early and late neonatal mortality rates were defined as proportion of dead neonates in first 7 and 28 days respectively. LBW, VLBW and ELBW were defined as birth weight under 2500, 1500 and 100, respectively. During 37-34 weeks of pregnancy, 34-30 weeks of pregnancy, and under 30 weeks of pregnancy, we consider PTB late, early, or very early [17]. Also, we can say, each predictor variable was examined separately for delivery to discharge survival for each patient, the most important of which were GA, neonatal birth weight, sex, and Apgar score. GA was calculated from the last menstrual period and sonography in the first half of pregnancy. In this paper, four models are studied for predicting the status of infant in the next sections.

## 3. Machine Learning Models/AI-General

There are many classification models and algorithms for predicting some parameters in the supervised ML/AI models. About classification models, some of important algorithms include “Logistic Regression”, “Decision Tree”, “Random Forest”, “Gradient-Boosted Tree, “XGBClassifier”, Multilayer Perceptron”, “Artificial Neural Network/MLP, “Gaussian Naive Bayes”, “K-Nearest Neighbors”, “Support Vector Machines”. The general road map and problem solution steps is depicted in a flowchart (algorithm) as an overall methodology which briefly is shown in Fig. 1.

**Fig. 1.**
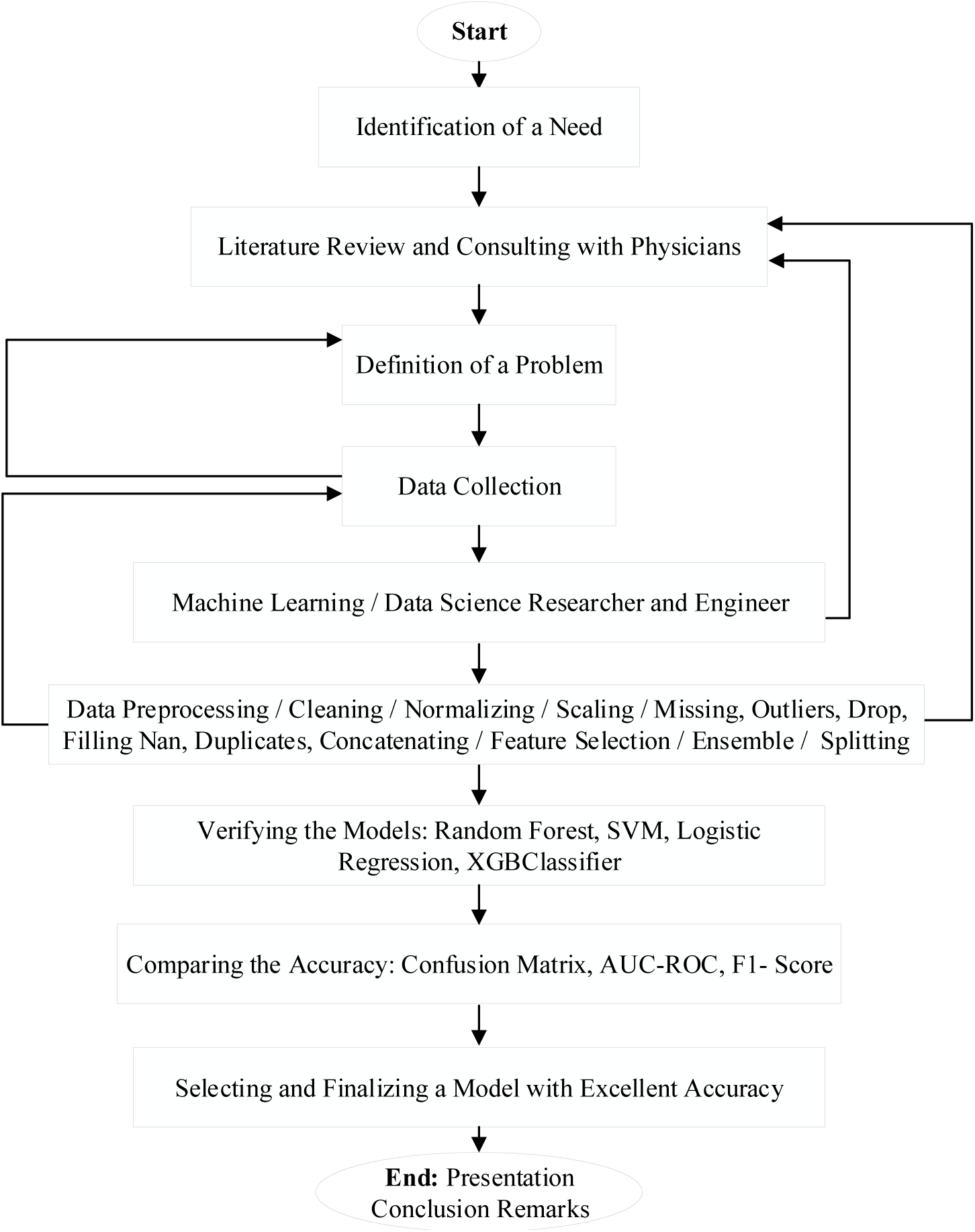
General road map and problem solution steps in a flowchart (algorithm): overall method

In this research paper, it has been tried to build and design a model with four ML/AI algorithms including Logistic Regression LR, Support Vector Machine SVM, and Random Forest RF, and XGBClassifier. Here, briefly these models are illustrated.

### 3.1. Logistic Regression

The initial and classic model of logistic regression (LR) is an instance of the supervised learning (SL). LR is used to compute and predict the probability of a binary (0/1 or yes/no) event occurring. A simple example of logistic regression can be employing in machine learning (ML) to obtain if a person is likely to be infected with a specific virus or not (see Fig. 2).

**Fig. 2.**
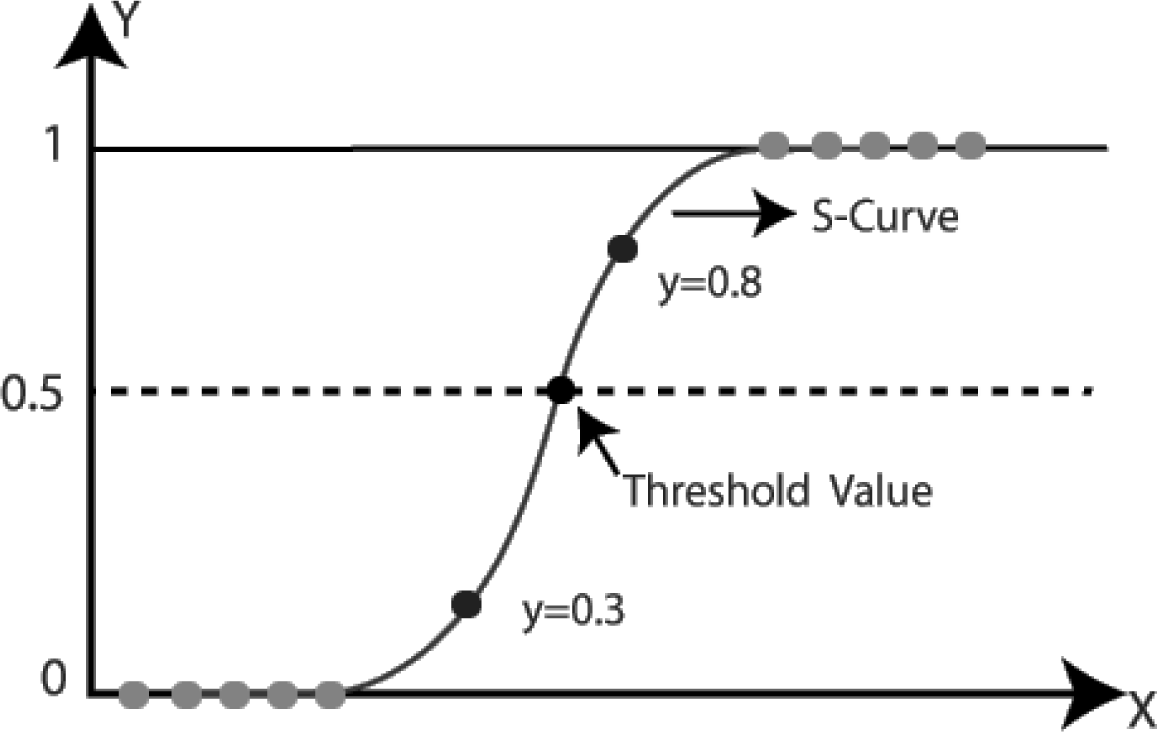
Logistic regression model [18]

Simply, the logistic function formulation is of the following Sigmoid form,

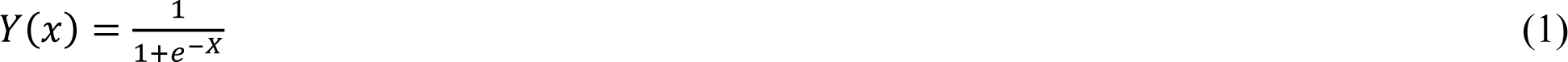

In which, logistic regression transforms its output utilizing the logistic sigmoid function to return a probability value generally. That is, LR is a special case of linear regression as it predicts the probabilities of result employing log-function. We apply the activation function (sigmoid) to change the outcome into categorical and classified values.

### 3.2. Support Vector Machine SVM

A support vector machine (SVM) is a supervised machine learning algorithm which employs classification algorithms for two group classification problems. SVM utilizes a method and technique called the Kernel trick to convert and transform a give data and so based on these transformations it finds a special optimum boundary between the possible outputs logically. Fig. 3 shows wholly components and terms of a SVM model schematically.

**Fig. 3.**
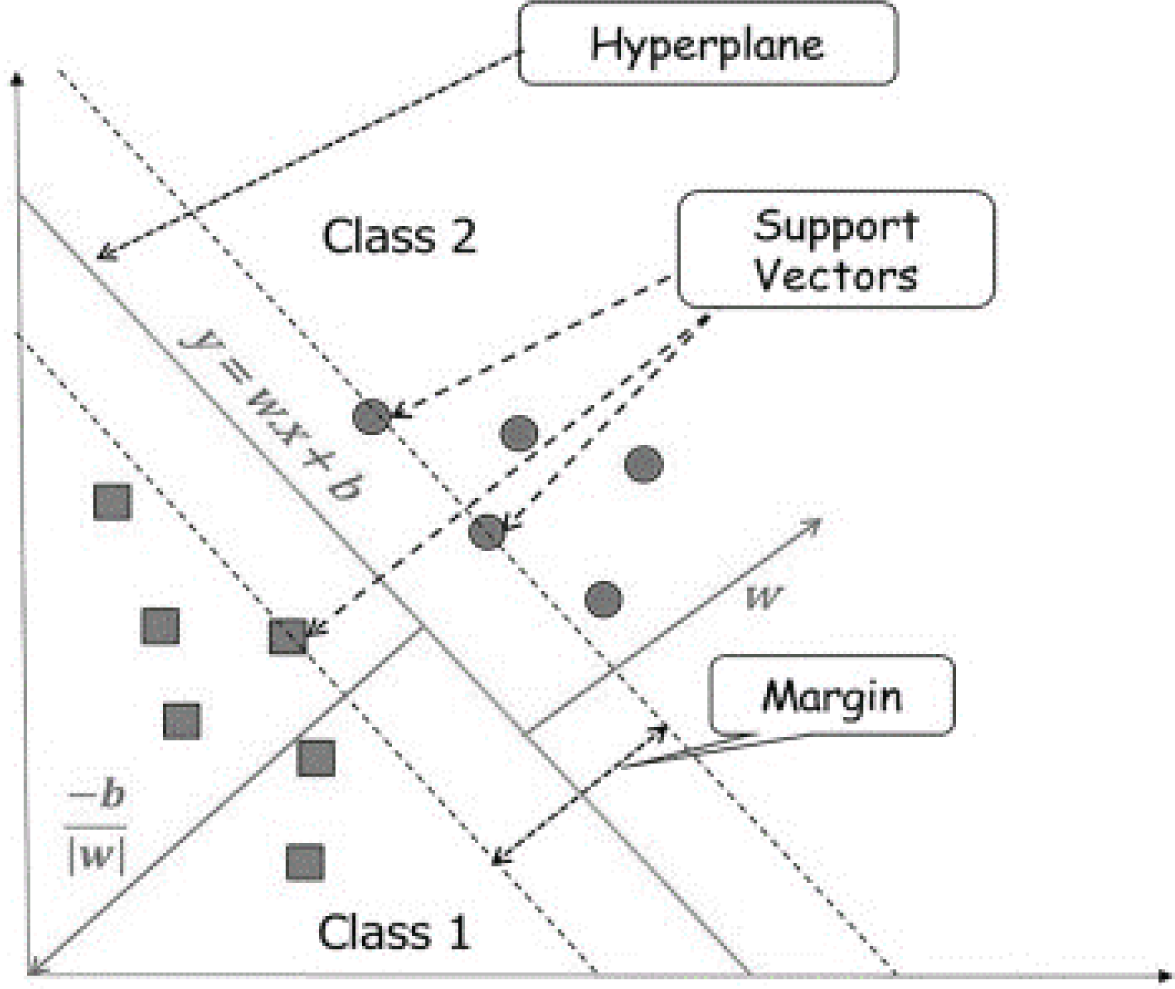
SVM model [19]

Regarding formulation of any hyperplane, it can be written as the set of points X satisfying the following formulation,

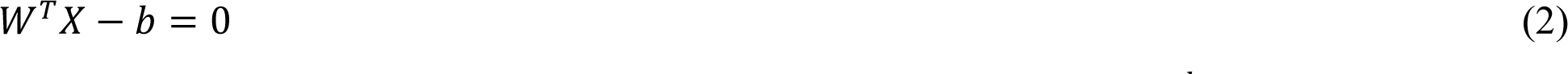

At which, W is the normal vector to the hyperplane and, the parameter 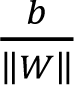 determines the offset of the hyperplane from the origin along the normal vector W.

### 3.3. Random Forest

The model of random forest is a supervised machine learning technique which is used extensively in classification and regression problems. It builds decision trees on dissimilar patterns and takes their majority vote for classification and average in case of regression. It should be mentioned that the model of random forests or random decision forests is an ensemble learning method for classification, regression and other tasks that operates by constructing a multitude of decision trees at training time. For classification tasks, the output of the random forest is the class selected by most trees generally (Fig. 4).

**Fig. 4.**
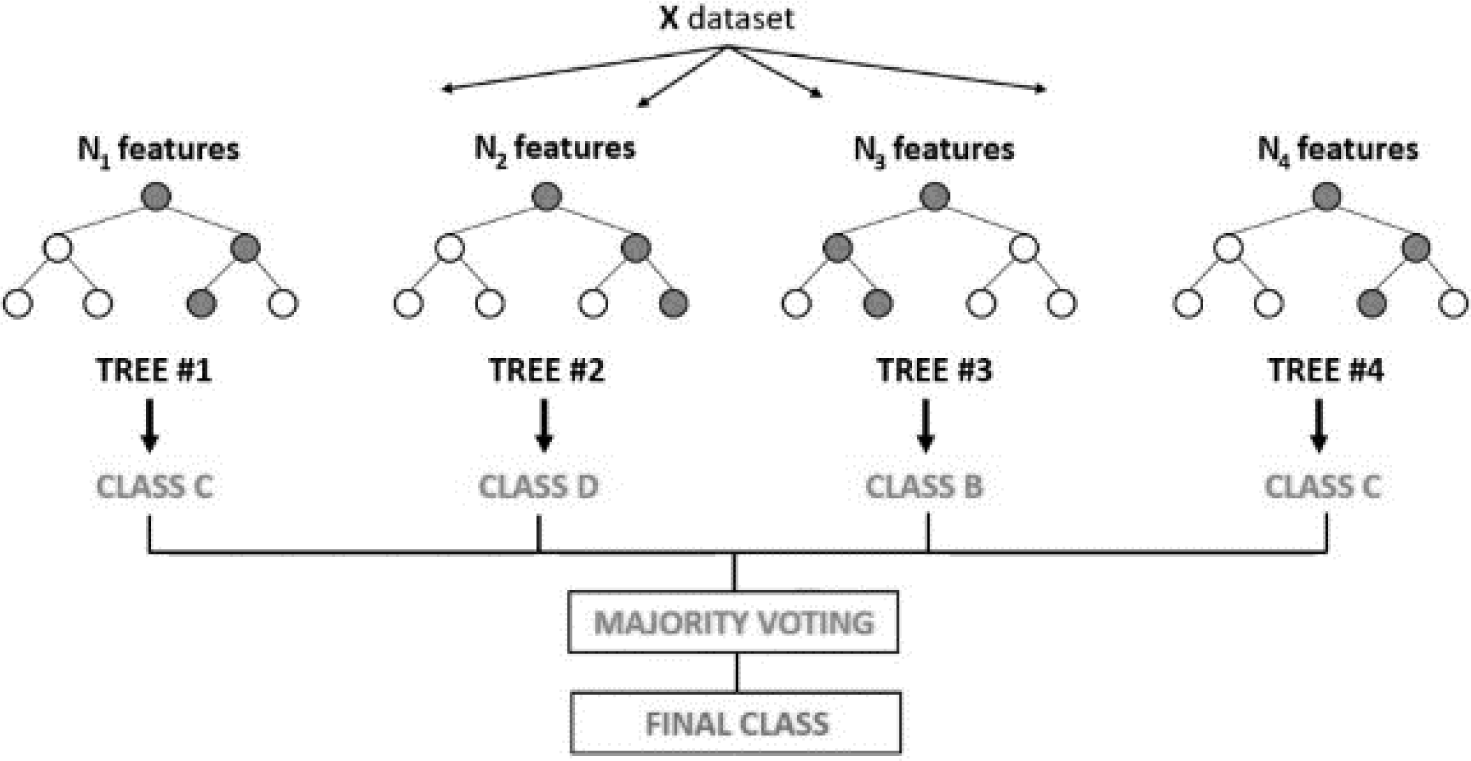
Random forest classifier model [20]

The solution steps for random forest were included choice random patterns from a given dataset and construct a decision tree for each pattern and sample and then get a prediction result from each decision tree. Then, doing a vote for each predicted result. Finally, select the prediction result with the most votes as the final prediction.

The mathematical meanings behind the random forest (∼ decision tree) consist of two important concepts: Entropy and Gain information. Entropy is a measure of the randomness of a system. Here, these two important formulations are presented as the following,

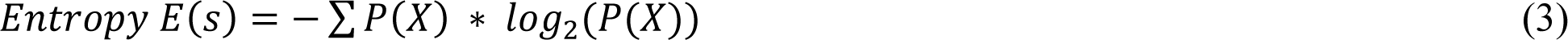

In which, P(X) is related to probability of X. Also, the Mathematical formulation for Entropy is as the following,

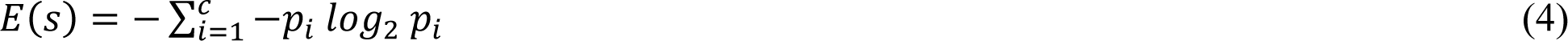

Where, c is number of classes, and 𝑝_𝑖_, is basically the frequentist probability of an element/class ‘*i*’ in our given dataset (𝑝_𝑖_is a fraction of examples in a given class). Also, regarding Gain information of Gain (S, A) have,

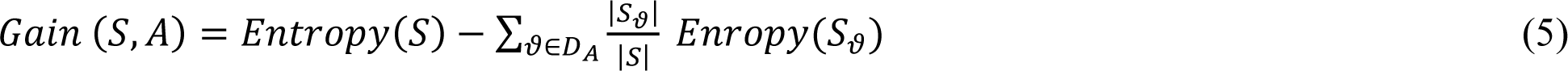

As well as, we have the following expression for Information Gain from X on Y,

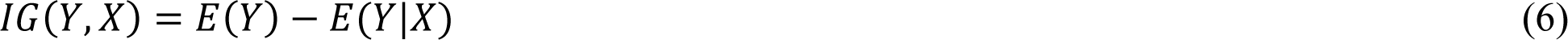

In which, we can subtract the entropy of Y given X from the entropy of Y to compute the decrease of uncertainty about Y given an additional part of information X about Y. Generally, it is called Information Gain “IG”. The larger the decrease in this uncertainty, the more information is obtained about Y from X logically.

Finally, the final vote and results is obtained by,

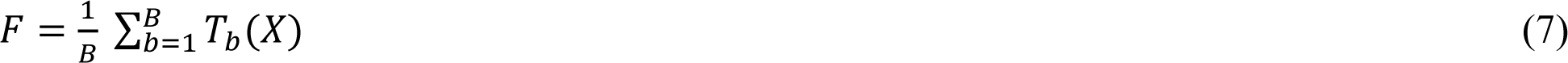

That is, after training, predictions for unobserved samples X may be made by averaging the predictions from all the individual regression trees on X field. The number of samples/trees, B, is a free parameter. Classically, several thousand trees are typically used, depending on the size and nature of the training set. An optimal number of trees B may be found utilizing cross-validation, or by detecting the out-of-bag error (bagging frequently: B times).

### 3.4. eXtreme Gradient Boosting (XGBoost)

eXtreme Gradient Boosting XGB is approximately more precise than random forests and more influential and stronger model. It combines a random forest and gradient boosting to build a stronger model and set of outcomes. XGB (XGBoost) has smaller steps, predicting sequentially instead of individualistically. It uses the samples and patterns in residuals, strengthening the model. It means the predicted error is less than random forest predictions.

In Extreme Gradient Boosting XGB model, Gradient boosting GB refers to a class of ensemble machine learning algorithms which may can be employed for classification or regression predictive modeling problems. Also, Ensembles are built from decision tree models generally (Fig. 5).

**Fig. 5.**
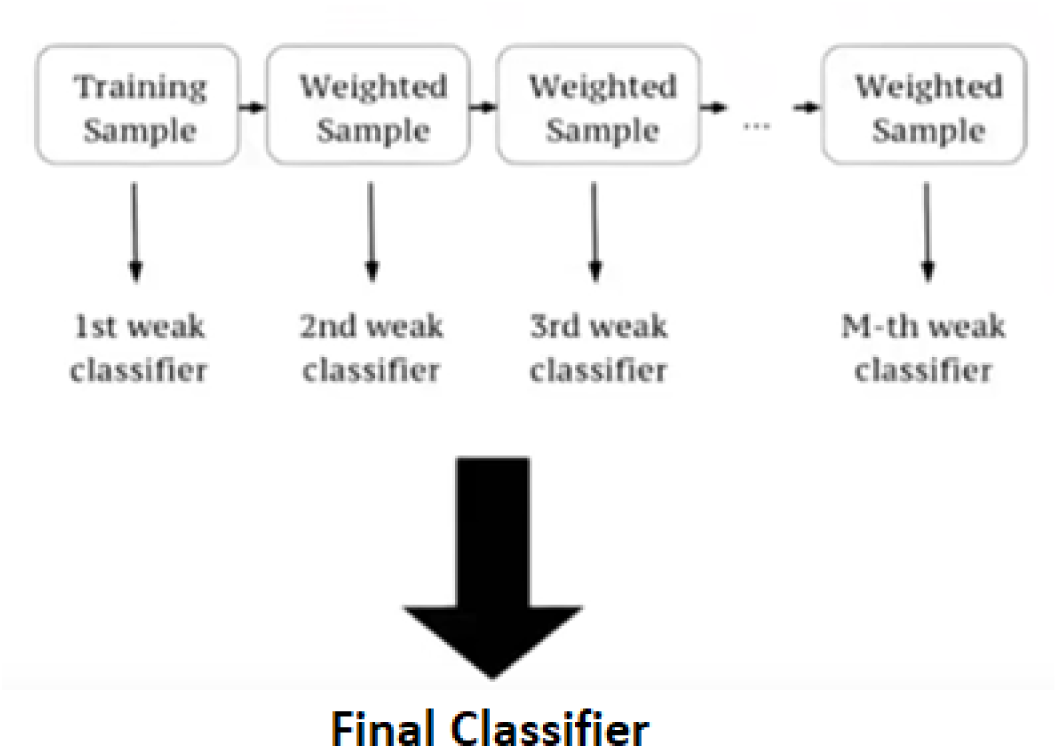
A simple model of XGB graphically [22]

XGBoost model works as Newton-Raphson in function field and space unlike gradient boosting which works as gradient descent in function space, a second order Taylor estimation is applied in the loss function to build the connection to Newton Raphson method. So, a general unregularized XGBoost algorithm is as follows. In XGBoost, we describe the complexity as below,

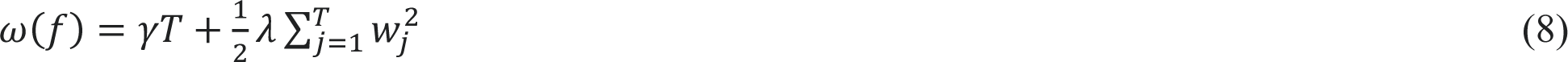

Where, 𝑤 is the vector of scores on leaves, *T* is the number of leaves, and 𝜆 is a regularization term.

## 4. Accuracy

There are two key kinds of errors current in any machine learning model. They are Reducible Errors and Irreducible Errors. The accuracy is a measure of the degree of closeness of a measured or calculated value to its actual value. The percent error is the ratio of the error to the actual value multiplied by 100. The precision of a measurement is a measure of the reproducibility of a set of measurements. Here, it is tried to explain briefly three types of accuracy (F1-Score, AUC-ROC, and Confusion Matrix). In this research, it is tried to use F1-Score for assessing the accuracy of the models.

### 4.1. F1-Score

F1-score is combining and balancing precision and recall on the positive class whereas accuracy looks at appropriately classified observations both positive and negative. F1 score is a generally utilized metric for classification machine learning algorithms, but its definition is not widely understood which can make it difficult know what a good score actually is. The formula used for F1-Score in is as follows,

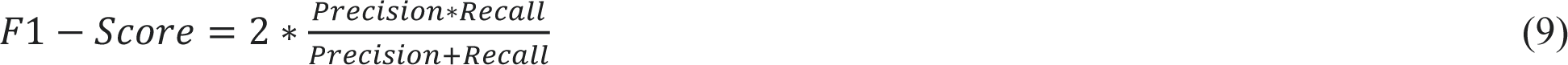

Precision is a quantity of how many of the positive predictions made are right (TP: True Positives), and have,

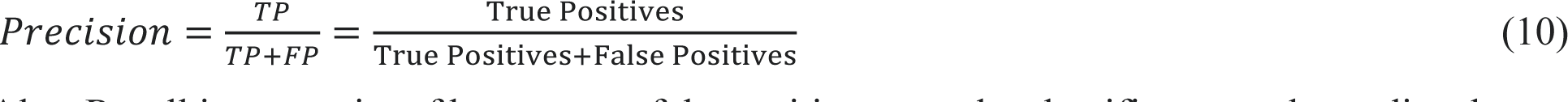

Also, Recall is a quantity of how many of the positive cases the classifier properly predicted, over all the positive cases in the given dataset, and defined as,

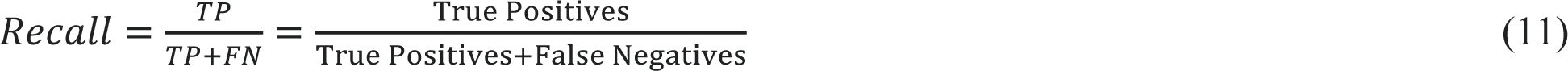

### 4.2. AUC-ROC

AUC-ROC diagram (curve) is a performance measurement for the classification problems at numerous threshold settings graphically. ROC is a probability curve, also and AUC presents the degree or measure of separability. It tells how much the model is capable of distinguishing between given classes. In ML (Machine Learning), performance measurement is a vital task. So, when it comes to a classification problem, we can count on an AUC - ROC Curve (Fig. 6). When we require to check or visualize the performance of the multi-class classification problem, we apply the AUC (Area Under the Curve) ROC (Receiver Operating Characteristics) curve. It is one of the most important evaluation metrics for checking any classification model’s performance. It is also written as AUROC (Area Under the Receiver Operating Characteristics) [22].

**Fig. 6.**
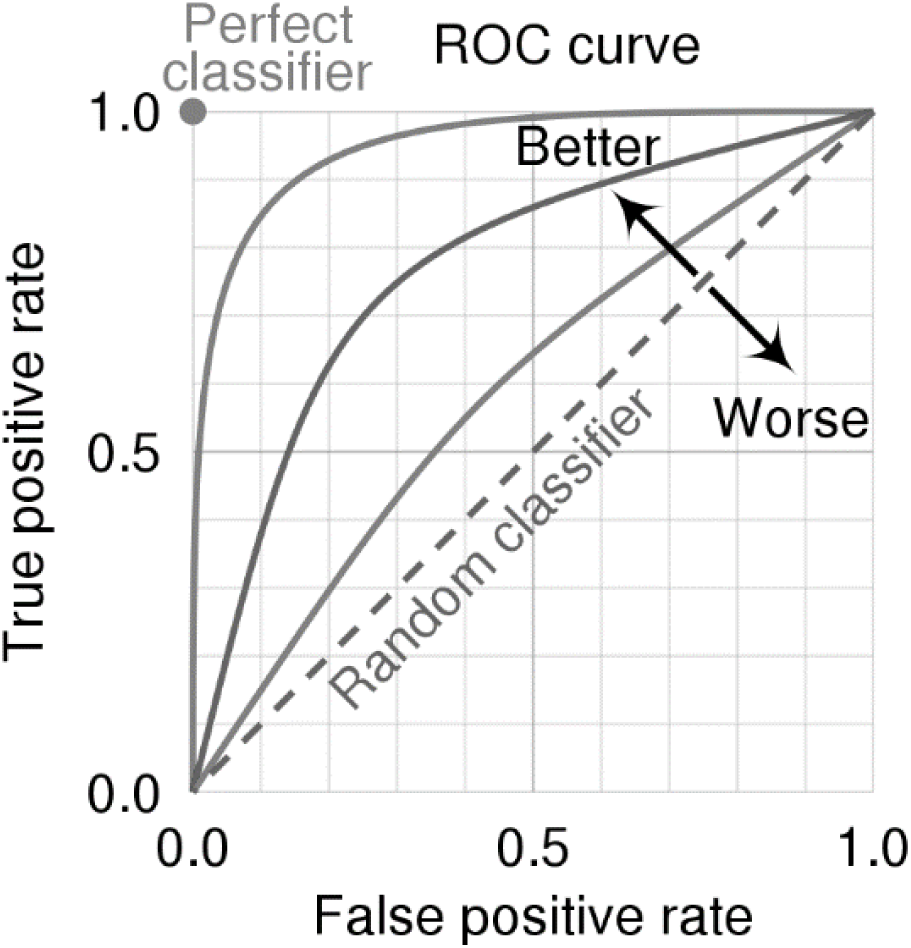
Presenting ROC-AUC curves and diagrams graphically

**Fig. 7.**
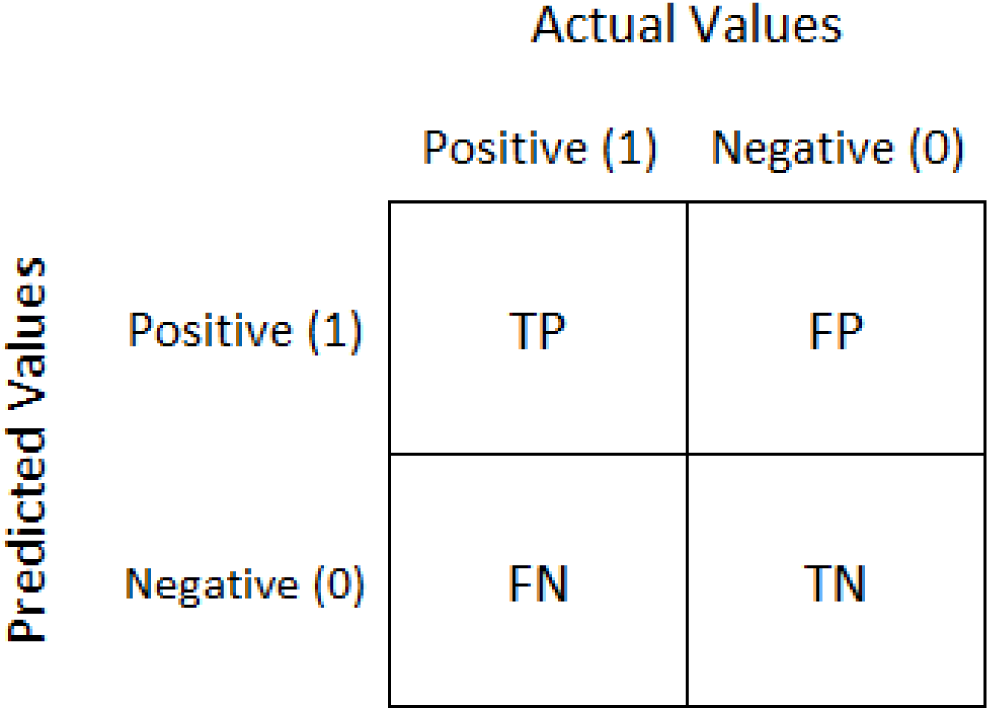
A simple sample of a confusion matrix schematically [23]

An AUC of 0.5 proposes no discrimination (capability to diagnose patients with and without the disease or condition based on the test), the values between 0.7 to 0.8 is assumed acceptable, and between 0.8 to 0.9 is considered excellent, and more than 0.9 is considered outstanding and ideal logically.

### 4.3. Confusion Matrix

In general, a confusion matrix CM is a numerical table which is applied to express the performance of a classification algorithm. A confusion matrix visualizes and summarizes the performance of a classification algorithm. The confusion matrices are employed to imagine and visualize significant predictive analytics like recall, specificity, accuracy, and precision. Confusion matrices are beneficial for the reason that they give direct comparisons of values like True Positives TP, False Positives FP, True Negatives TN and False Negatives FN.

Confusion matrices are extensively used because they give a better concept of a model’s performance than classification accuracy does. For example, in classification accuracy, there is no information about the number of misclassified instances. The confusion matrix is denoted by a positive and a negative label and class. The positive label denotes the not-normal class/label or behavior, so it is typically less represented than the other label/class. The negative label, alternatively, represents normality or a normal behavior.

## 5. Preprocessing the Dataset

Preprocessing the data is the heart of machine learning. Your machine learning instruments are as good as the quality of your data. our data requires to go through a few steps before it could be used for making predictions. Here, it is presented a short drawing concerning how to preprocess data in Python step-by-step as the following,

- Get the dataset and import all the crucial libraries
- Load and import data in Pandas
- Drop columns that aren’t useful (feature selection)
- handling data including removing outliers and duplications, dropping/filling NAN data
- Drop rows with missing values or substituting with mean, median or other techniques
- Feature scaling (standardization, normalization)
- Create dummy variables and cleaning data set
- Data transformation
- Data reduction and dimensionality reduction
- Encoding the categorical data
- Balancing the data
- Take care of missing data and screening
- Convert the data frame to NumPy
- Divide the data set into training data and test data (Splitting the dataset)

So, Preprocessing the dataset is essential and necessary to model an AI/ML algorithm for prediction of parameters. It means that preprocessing the dataset is done to get the important and effective features (feature selection) and increasing the accuracy. There are three usually employed feature selection methods that are simple to make and yield excellent results and good model as the following,

### A. Correlation Matrix with Heatmap

With this technique, we can see how the features are correlated with each other and the target. The Correlation Matrix shows Positive output if the feature is highly relevant and will show a Negative output if the feature is less relevant to the data. A Heatmap always makes it easy to see how much the data is correlated with each other and the target. In this way, we can select the most relevant features from your dataset using the Feature Selection Techniques in Machine Learning with Python (heatmap method).

### B. Feature Importance

With this technique, we can get the feature importance of every feature from your dataset with the use of feature importance tool of the model. Feature Importance works by giving a relevancy score to you to every feature of your dataset, the higher the score it will give, the higher relevant that feature will be for the training of your model (ExtraTreesClassifier model).

### C. Univariate Selection

Statistics may be utilized in the selection of those features that carry a high relevance with the output. In this paper, the statistical test method is used for the positive features to select the best features from the dataset (chi2).

### D. Logistic Regression

We may fit a Logistic Regression model on the regression dataset and recover the Coeff_ property that includes the coefficients found for each input variable and feature. These coefficients can give the basis for a common feature importance score. This presumes that the input variables have the same scale or have been scaled before fitting a model.

### E. XGBoost Feature Importance

XGBoost is an important library which suggests a capable application of the stochastic gradient boosting SGB algorithm. This algorithm may be utilized with scikit-learn via the XGBRegressor and XGBClassifier classes usually. Also, this model and algorithm is presented through scikit-learn via the Gradient Boosting Classifier and Gradient Boosting Regressor classes and the same methodology to feature importance and selection may be employed.

<u>Note</u>: According to Fig. 1, cleaning the dataset is required including management and optimizing the outliers, missing values, balancing the data, normalization, duplicated data, number of data and predictors/attributes, and error in collecting data.

## 6. Results and Discussions

### 6.1. Summary of Method

After building models and implementing numerous machine learning algorithms on the mentioned features, we found that the better results are determined for four mentioned models developed above. Many models have been initially developed by means of the original data; but, because of the low sample in “dead” class, the model performance is not acceptable logically. So, to get the best models, data pre-processing techniques are necessary and must be applied for balancing, tuning, adjusting the data, as well as cleaning the data (standardization of dataset for starting the learning). Also, data imputation is typically applied to improve the dataset quality and help to machine learning models for better performance. To get this, many models and algorithms are well-known and documented like replacing the average/mean/median or mode of a class group. In ML/AI “SMOTE” is the most well-known method to balance the data to improve random oversampling as well as logically setting the dataset (Fig. 8).

**Fig. 8.**
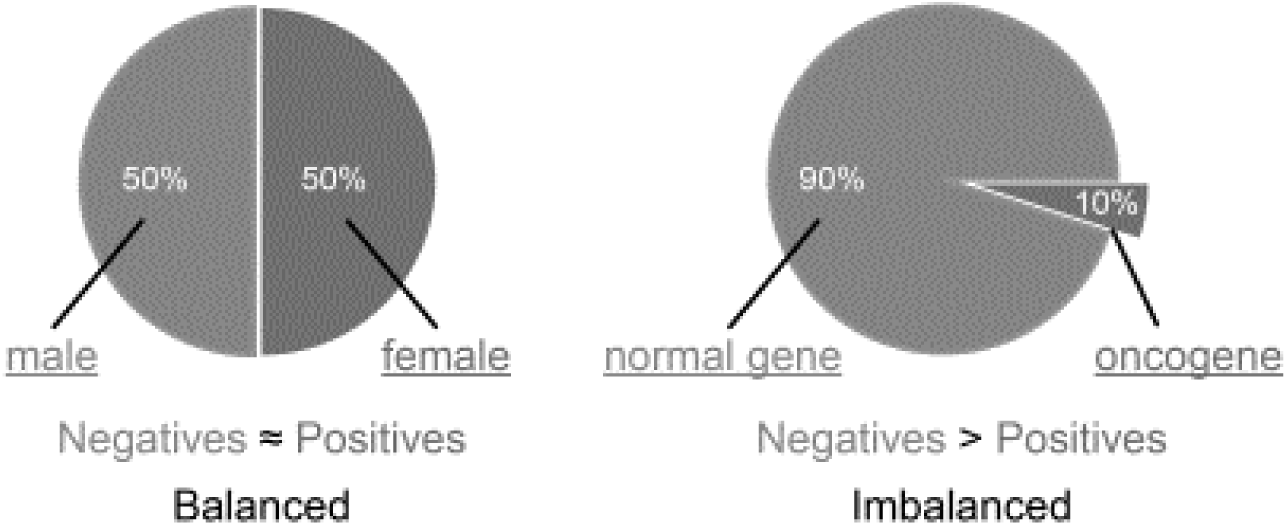
Graphical example of balanced and imbalanced data [24]

Based on the pre-processing methods, we built a new dataset using Python software. These methods are including data preprocessing, cleaning, normalizing, scaling, standardizing, revising the missing, outliers, Nan, and duplicates data, dropping, filling Nan, feature selection, ensemble, and splitting. Sometimes, feature selection may be useful for getting a suitable accuracy. To get the best outcomes, different machine learning algorithms were implemented on the novel data, and the above-mentioned oversampled datasets and compared the performance of the algorithms in terms of confusion matrix measures.

After preprocessing the datasets, the variables and features as input variables are used to develop the machine learning models. The data were divided (split) into two major groups: 75% for training and 25% for testing data and then dissimilar machine learning algorithms including SVM, random forest, XGB, and logistic regression were developed on the original and the oversampled datasets. Also, each algorithm was done 20 times with different randomly selected train and test sets.

### 6.2. Detailed Results

Here, some important results and outcomes are shown after implementing the ML/AI models in Python. It is necessary that indicated to assumption of digit “1” for our aim (dead) as positive. In this paper, it has been tried to compare three methods of SVM, random forest and logistic regression results and accuracies. First, heat map results are depicted and shown for more clarification of the effects of the parameters (features) and their significances graphically as the following (Fig. 9). A heat map helps to visualize density. On the other hand, a heat map is a 2D representation of dataset at which values are represented by various colors. A simple heat map provides an immediate visual summary of information. More elaborate heat maps allow the viewer to understand complex data sets (Correlation Matrix with Heatmap).

**Fig. 9.**
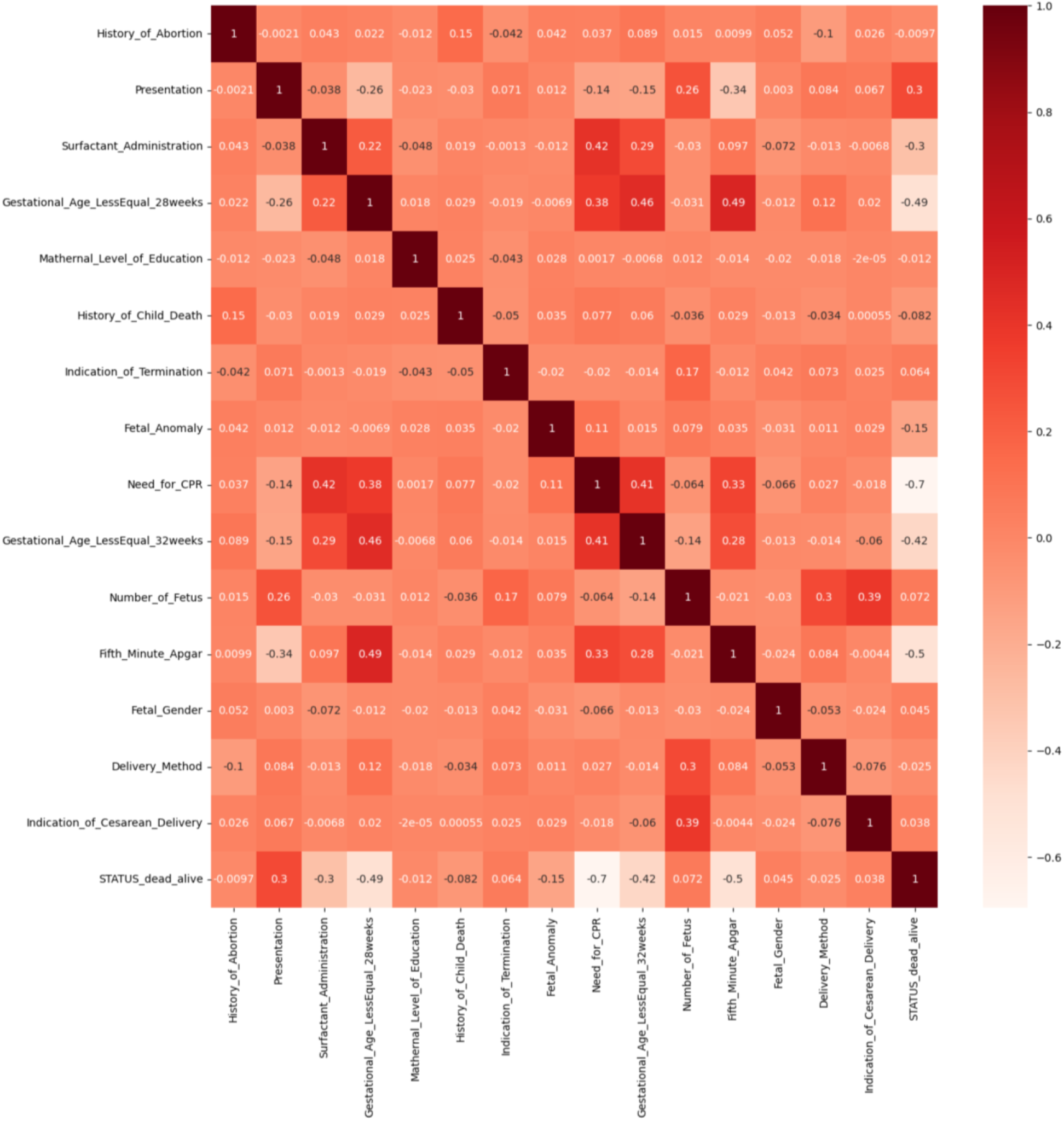
A heat map table for showing the importance of the features and their relations

All numbers between 0 to 1 are meaningful in this table and show the importance of the features. That is, the features with values of nearing 1 (dark color: colorful) have a maximum effect on the final result (dead or alive) unlike zero (light color: colorless) for features. In this figure, it found that “need for CPR” is the most important factor for surviving of the infant (−0.7).

Then, the second method results for feature selection would be “Feature Importance”. With this method, we may get the feature importance of every feature from your dataset with the use of feature importance tool of the model (Fig. 10). Feature Importance works by giving a relevancy score to you to every feature of your dataset, the higher the score it will give, the higher relevant that feature will be for the training of your model (Extra Trees Classifier model). The result of this method is as the following,

**Fig. 10.**
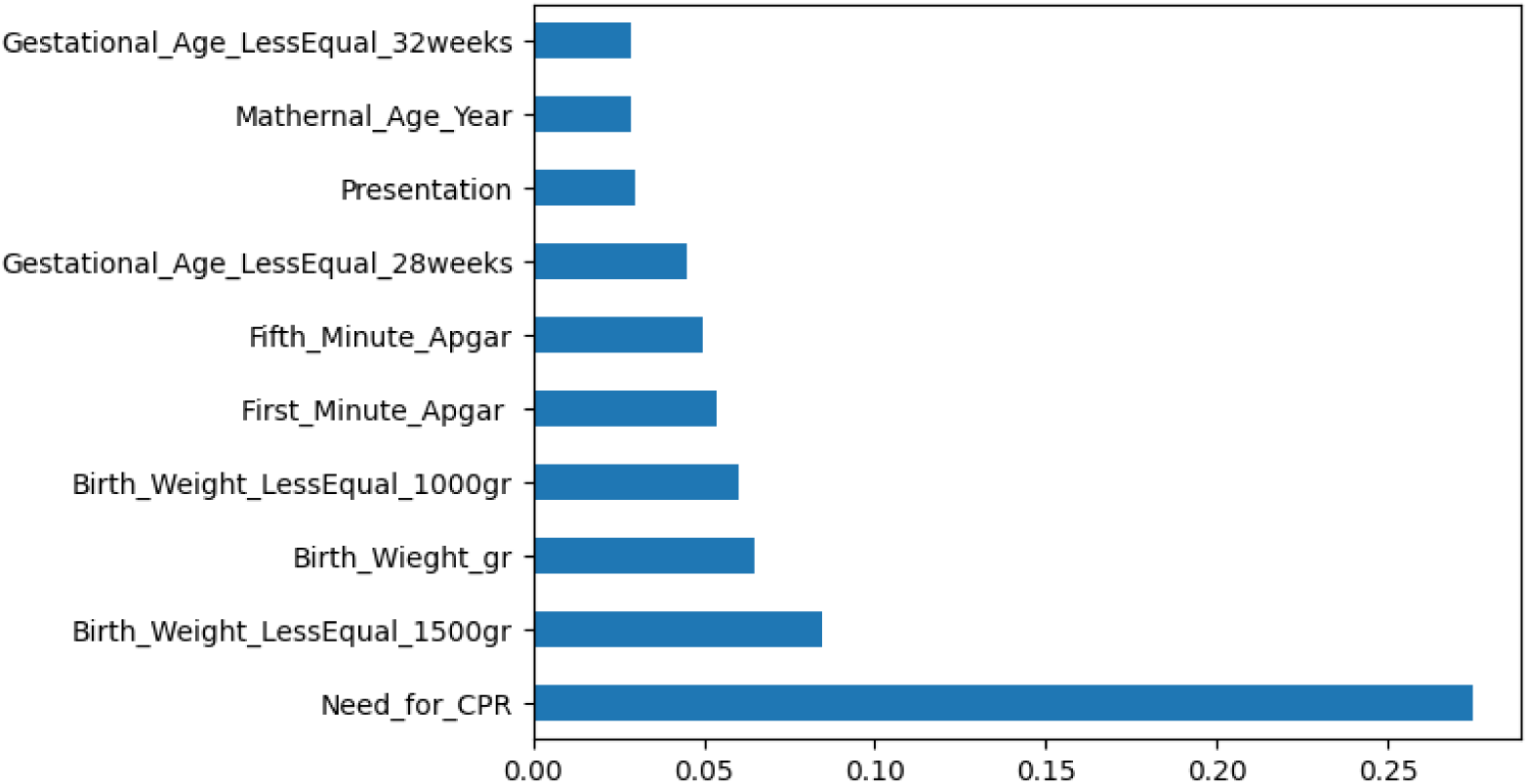
Bar chart for feature selection by Extra Trees Classifier model

In this figure, it found that “need for CPR” is the most important factor for surviving of the infant (∼ 0.28).

The third method is “Univariate Selection”. Statistics can be used in the selection of those features that carry a high relevance with the output. In this paper, the statistical test method is used for the positive features to select the best features from the dataset (chi2). The obtained result of this method is as the following (Fig. 11),

**Fig. 11.**
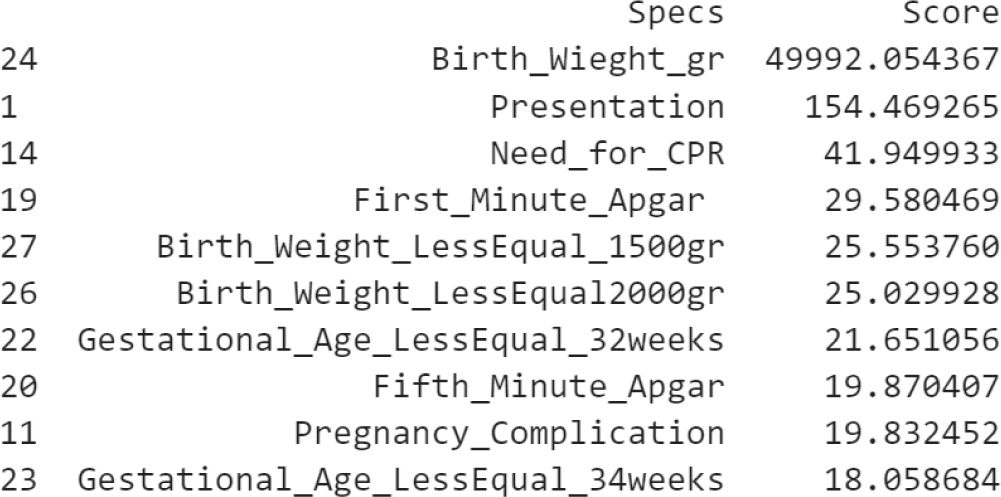
The table for feature selection by chi2 model

In this figure, it found that “Birth_Weight_gr” is the most important factor for surviving of the infant with score of 49992.054367.

The fourth method for feature importance is “Logistic Regression”. We may fit a Logistic Regression model on the regression dataset and recover the Coeff_ property that includes the coefficients found for each input variable and feature. These coefficients can give the basis for a common feature importance score. This presumes that the input variables have the same scale or have been scaled before fitting a model. The obtained result of this method is as the following (Fig. 12),

**Fig. 12.**
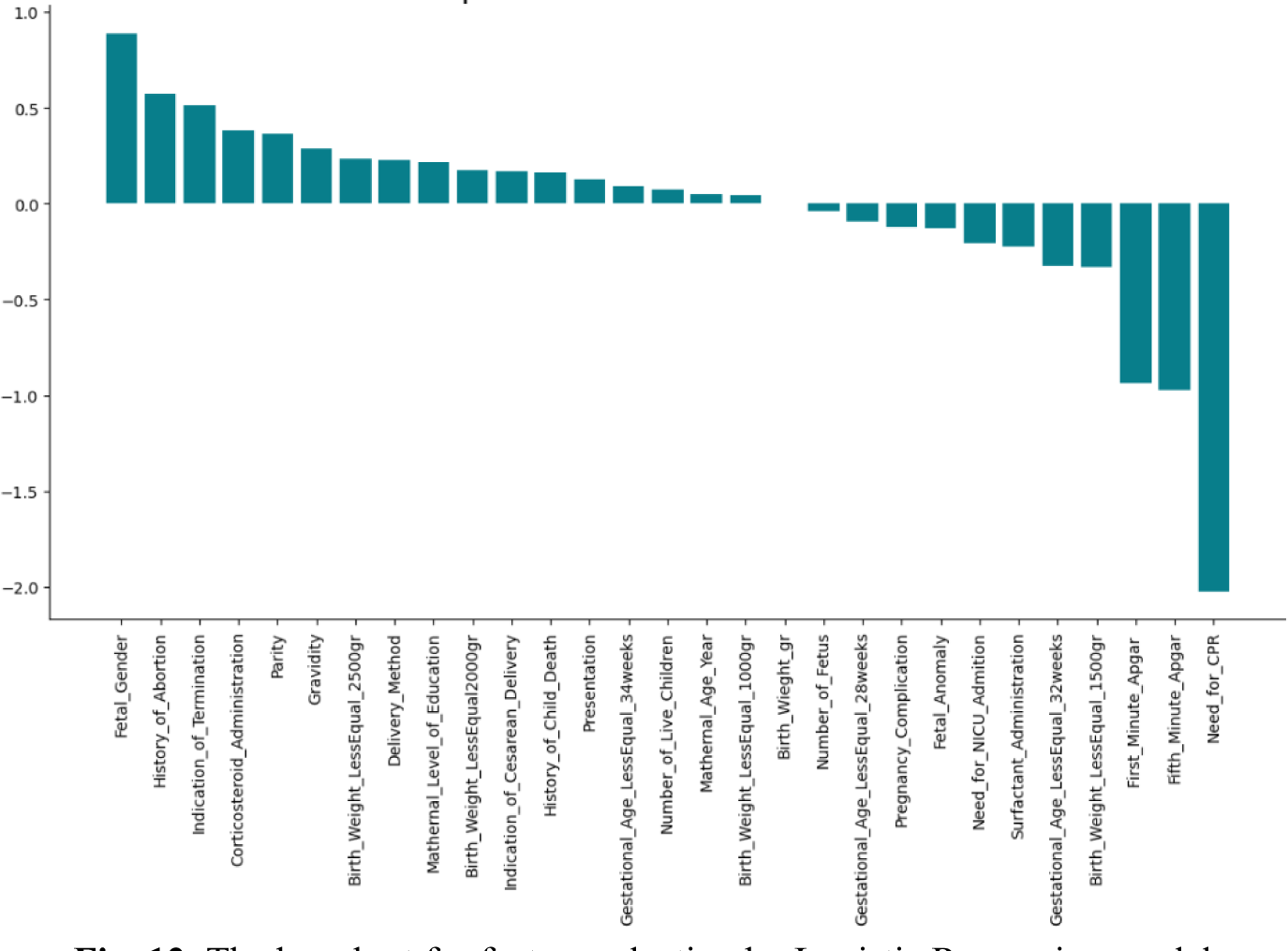
The bar chart for feature selection by Logistic Regression model

In this figure, it found that “Need_for_CPR” is the most significant factor for surviving of the infant with score of about -2.0.

The fifth method for feature importance is “XGBoost Feature Importance”. In which, we would have the following outcomes (Fig. 13),

**Fig. 13.**
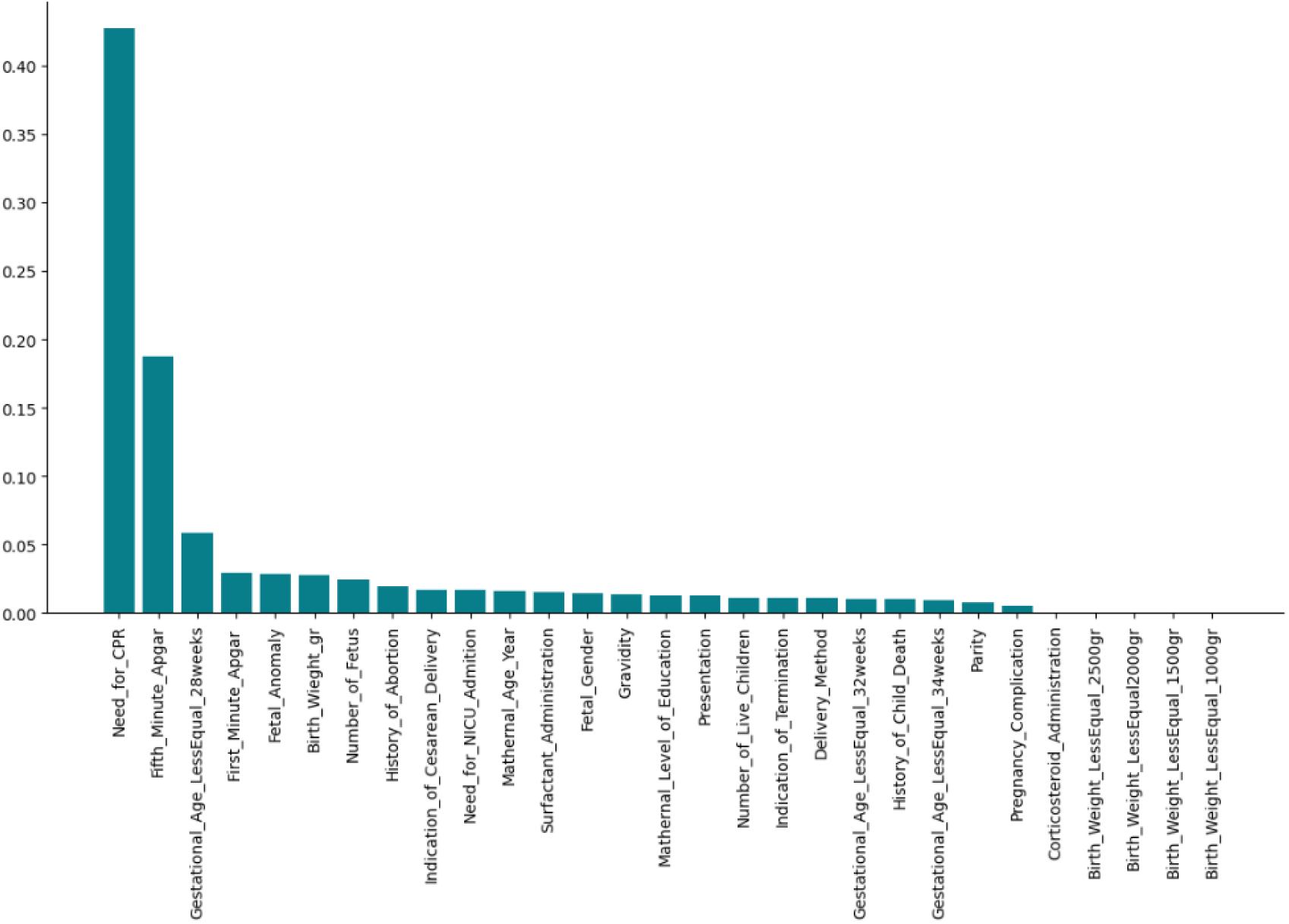
The bar chart for feature selection by XGB model

In this figure, it found that “Need_for_CPR” is the most significant factor for surviving of the inf ant with score of about ∼ 0.43. So, about 15∼20 important features can be found and extracted via the mentioned feature selection methods averagely. For example, union top features in XGB and Coeff methods are including ’Parity’, ’Surfactant_Administration’, ’Birth_Weight_LessEqual_150 0gr’, ’Gestational_Age_LessEqual_28weeks’, ’History_of_Child_Death’, ’Indication_of_Terminati on’, ’Fetal_Anomaly’, ’Number_of_Fetus’, ’Fifth_Minute_Apgar’, ’Birth_Wieght_gr’, ’Number_of_ Live_Children’, ’Gravidity’, ’Pregnancy_Complication’, ’Mathernal_Level_of_Education’, ’Birth_ Weight_LessEqual_2500gr’, ’Delivery_Method’, ’Indication_of_Cesarean_Delivery’, ’Need_for_C PR’, ’Gestational_Age_LessEqual_34weeks’, ’History_of_Abortion’, ’Presentation’, ’Fetal_Gender’, ’First_Minute_Apgar ’, ’Need_for_NICU_Admition’, ’Mathernal_Age_Year’, ’Gestational_Age_LessEqual_32weeks’, ’Birth_Weight_LessEqual_1000gr’. Also, intersection top features in XGB and Coeff methods are comprising ’History_of_Abortion’, ’Presentation’, ’Surfactant_Administrati on’, ’Gestational_Age_LessEqual_28weeks’, ’Mathernal_Level_of_Education’, ’History_of_Child_Death’, ’Indication_of_Termination’, ’Fetal_Anomaly’, ’Need_for_CPR’, ’Gestational_Age_Less Equal_32weeks’, ’Number_of_Fetus’, ’Fifth_Minute_Apgar’, ’Fetal_Gender’, ’Delivery_Method’, ’ Indication_of_Cesarean_Delivery’.

As a results, the any researcher or engineer can select one or more than one method for feature selection or combination and/or intersection of those as needed.

In addition to the mentioned methods, PCA (principal component analysis) is an available method which is approximately similar to the feature selection methods. Here, a result of PCA is shown graphically as below (Figs, 14, 15),

Principal Component Analysis (PCA) is a wonderful method for dimensionality reduction and may also be utilized to define feature importance. PCA won’t show us the most significant features at once, as the earlier mentioned techniques did. As a substitute, it will return N principal components, where N equals the number of original attributes and features (see Figs. 14,15). Here, PCA loading scores are presented for the first principal component pc1. In which, we can extend it from pc1 to pc29 or less.

**Fig. 14.**
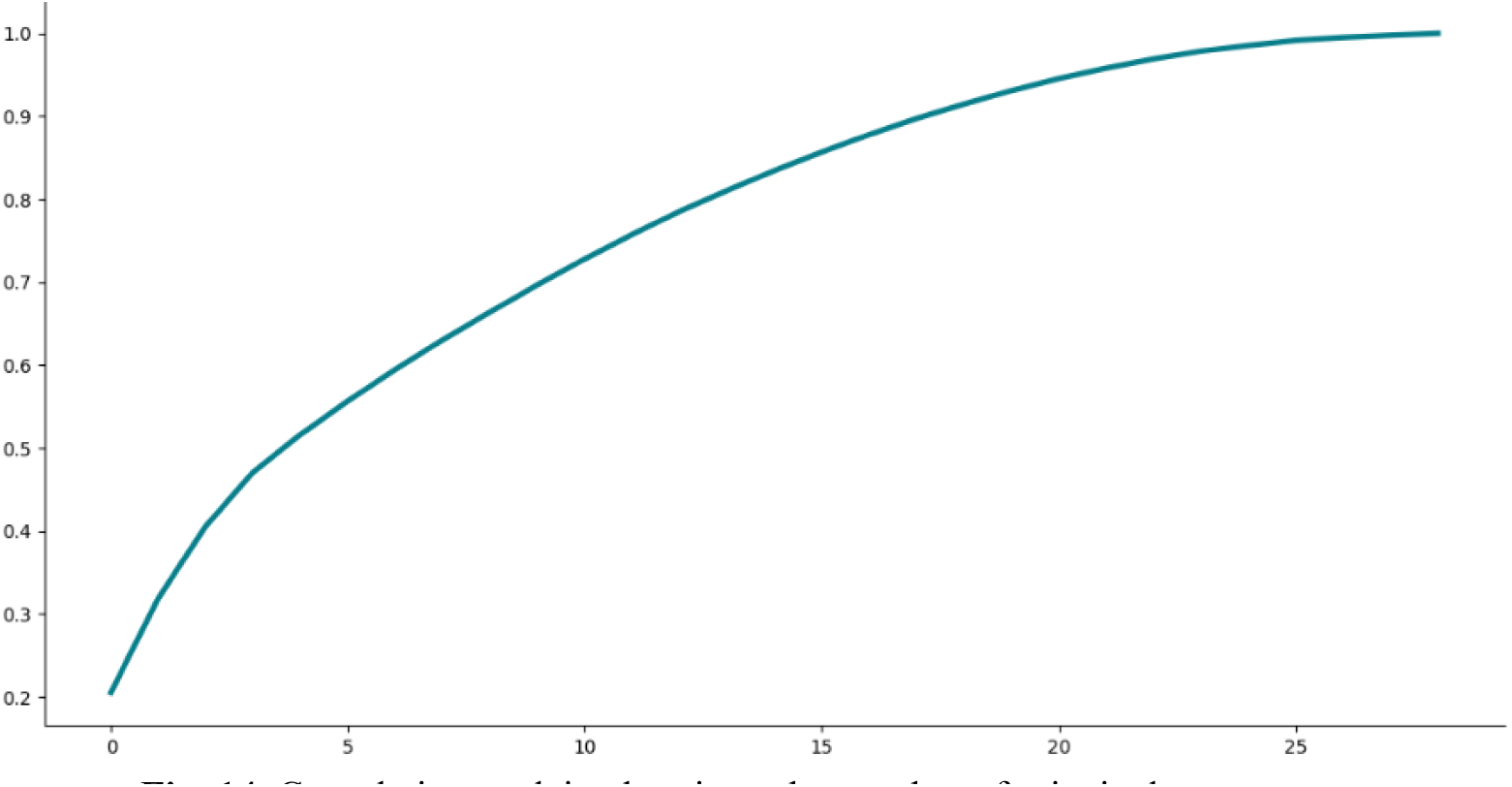
Cumulative explained variance by number of principal components

**Fig. 15.**
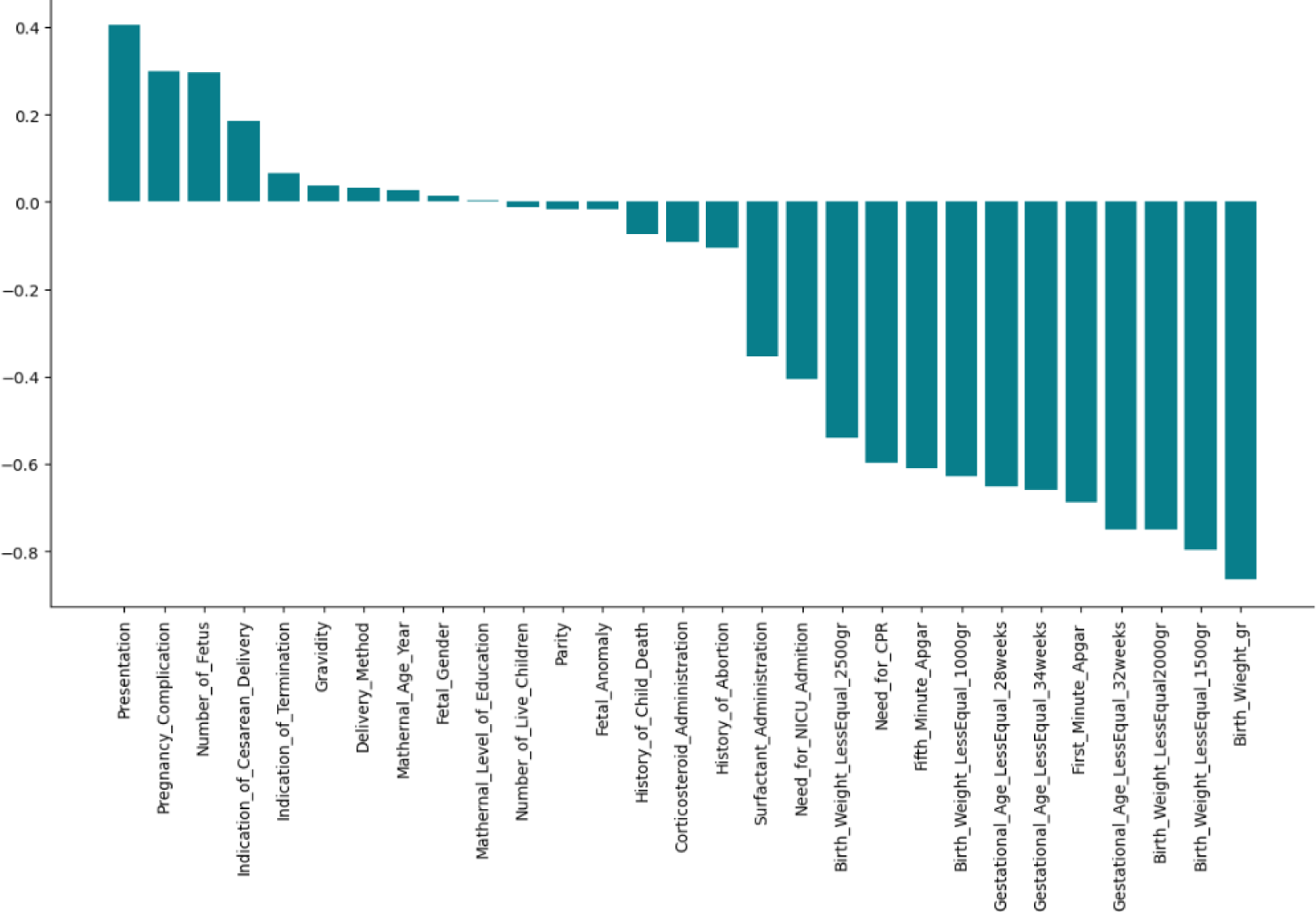
PCA loading scores (first principal component pc1)

In the following, the obtained results are presented for comparing the accuracy of the mentioned models as below (Table 3)

**Table 3.** Models and errors (average)

| No | Model | Accuracy (F1-Score), % |
| --- | --- | --- |
| 1 | Logistic Regression | 87.50 |
| 2 | Random Forest | 90.45 |
| 3 | Support Vector Machine SVM | 90.35 |
| 4 | XGBClassifier | 91.50 |

Table 3 shows that the optimum model with simplicity and suitable accuracy is Random Forest algorithm which has a low bias/error, and variance. As a result, CPR is one of effective features which affect directly on the infant health. For finding more feature which are significant, please refer to the related section above.

Generally, around 20 factors have been finalized which are significant in the neonatal mortality prediction. F1-Score was obtained for our three models and SVM and RF were suitable models for this problem. The best score (accuracy) was 90.45 for RF model. The highest accuracy (F1-score) is for RF model.

Also, a confusion matrix CM can help more to have a better vision of the problem. It is a numerical table applied to state the performance of a classification algorithm. Here, a result is presented for more clarification (Table 4). Confusion matrices are widely employed because they give a better idea of a model’s performance (with more details) than classification accuracy does.

**Table 4.**
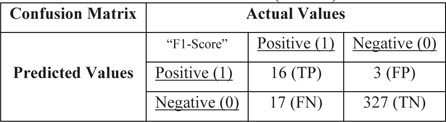
Confusion matrix (F1-score)

If we consider and assume the number of one (positive) for dead status and zero (negative) for alive status of infant after delivery, the obtained results would be interesting. In fact, for reminding, it should be mentioned that this table has been obtained for test datasets. It means that 𝑇𝑃 = 16, 𝐹𝑃 = 3, 𝐹𝑁 = 17, and 𝑇𝑁 = 327. The main challenge is in value of 𝐹𝑁 (= 17), and we must design and build an AI/ML model to decrease it. It means that the AI model has wrongly predicted the status of infant (these 5 infants have died or would be died while AI model has predicted they are alive or would be alive/survived). Here, 𝐹𝑁 is a critical factor that must be optimized by AI model.

As one of a real table, here it is shown graphically as below (Table 5),

**Table 5.**
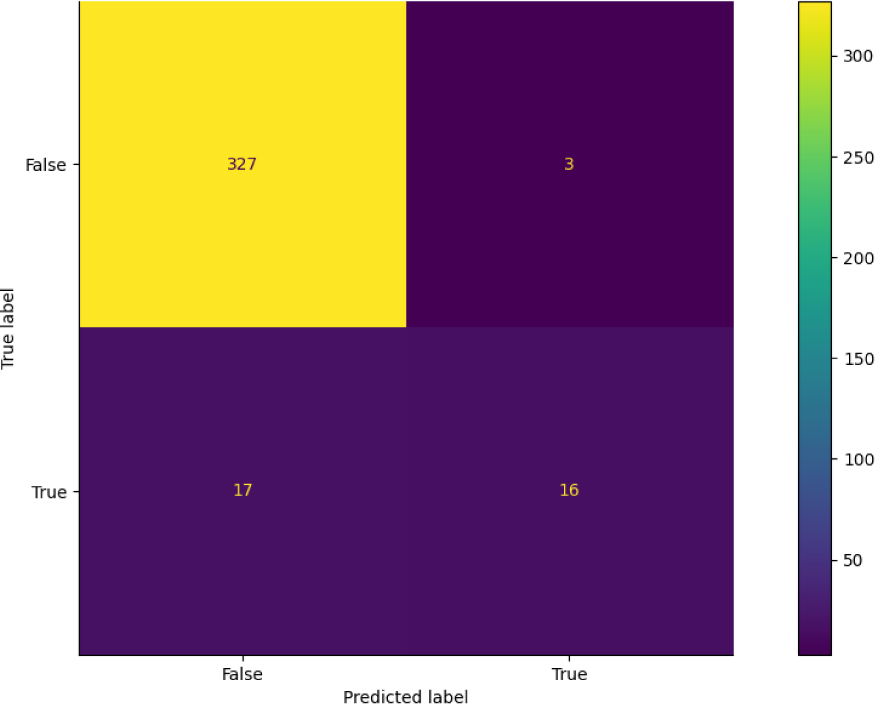
Confusion matrix (F1-score) before optimization.

Table 5 (TP=16, TN=327, FP=3, FN=17) is an initial confusion matrix for one of our model. As shown, 𝐹𝑁 = 17, so we must try to decresae it by some process like selection of model, features, tuning and setting some parameters, and what have you. These processes have been described before (in this text) comprehensively. That is, we have to design another model to decrese this FN. So, it is neede to redesign and rebuild another efficient AI model which will be done in then next paper. Note that, in this classificaion problem, The dead class is underrepresented (not-normal) class wherein the alive class is the normal class in this data set. Therefore, the dead class would represent the positive class, while the alive class would represent the negative class in the confusion matrix (the class/label with less samples would be positive).

As a final diagram, ROC curve is depicted for more visualization. ROC curve is a chart that visualizes the tradeoff between true positive rate TPR and false positive rate FPR. Mainly, for every threshold, it is better to estimate TPR and FPR and plot it on one chart. For sure, the higher TPR and the lower FPR is for each threshold the better and so classifiers that have curves that are more top-left side are better logically (Fig. 16).

**Fig. 16.**
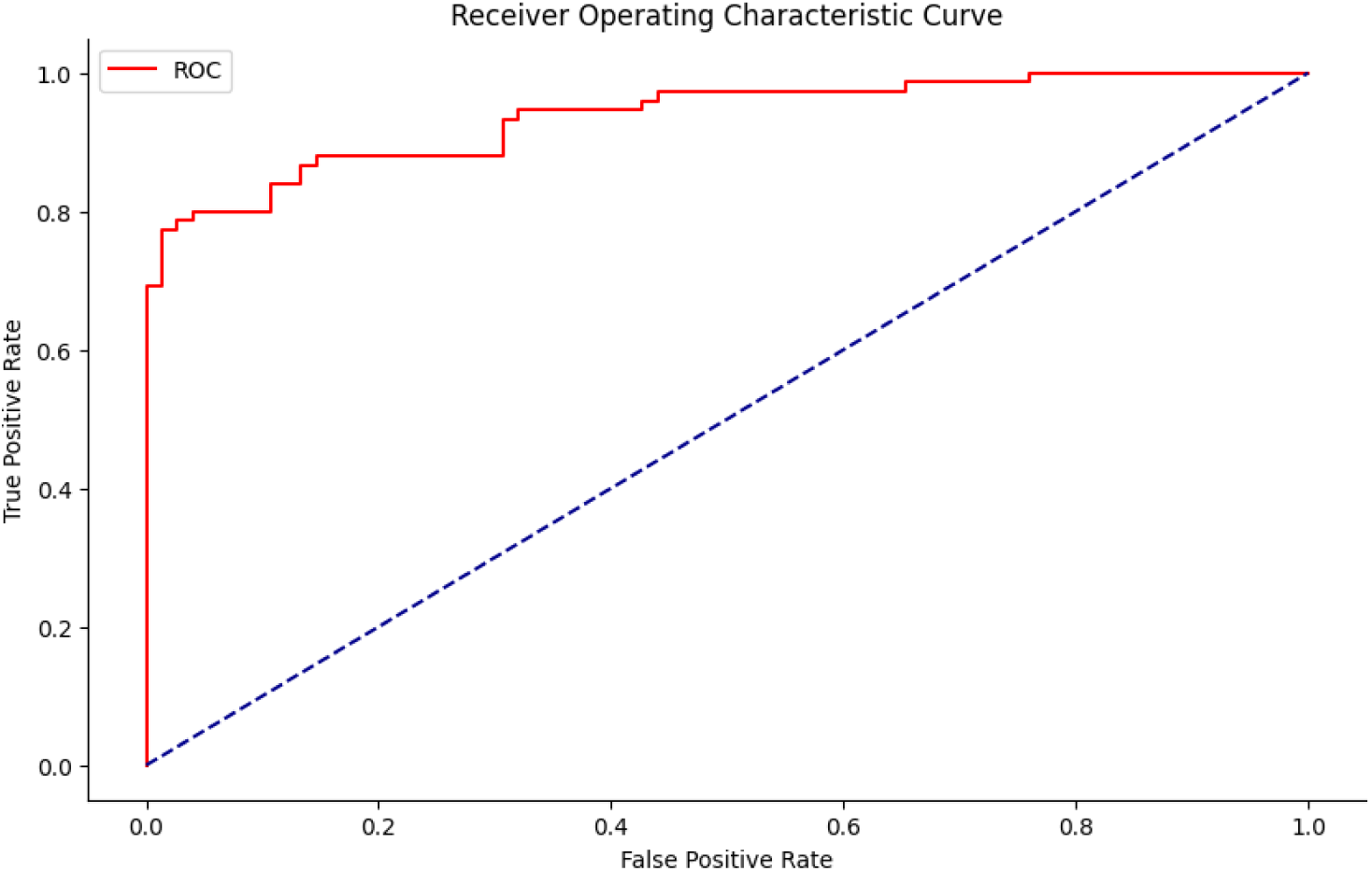
ROC curve for this project

The area under red curve is approximately near one, that is, the model has an acceptable and suitable accuracy (Fig. 16). Finally, after learning and selecting a suitable model, the physicians or any hospital personnel can predict the status of infant (dead or alive) with excellent accuracy (Table 3) in accordance with the mentioned obtained important features (∼ 20 features). That is, for instance, the simple Python code of “status_dead_alive = model.predict(new patient conditions or features)” can predict the status of infant (dead:1 or alive:0) with high accuracy.

## Conclusion

Different feature selection methods were used to develop an ensemble of machine learning-based models to predict neonatal deaths in NICUs by Logistic Regression LR, Random Forest RF, Support Vector Machine SVM, XGBClassifier, and Ensemble methods. In consequence, models were developed based on features selected by neonatologists. Also, use of heat-map is proposed for having better vision and suitable imaging of effects of the features on the output to better designing the model. In prospective evaluations, XGBClassifier model performed best. As a result, it is recommended for use on similar projects. In addition to clean, real and valuable dataset, an excellent model was built and designed for predicting the final health status of the infants at the hospital. So, with using this algorithm, the physicians can do any required caring for surviving for the infants with high probability of dying. As a result, CPR is one of efficient features which affect directly on the infant health (for finding more important features please refer to the results). In a word, employing the developed artificial intelligence/machine learning (ML/AI) algorithms may assist and help physicians to predict the neonatal deaths in NICUs with high accuracy.

## References

[1] Repka MX. Ophthalmological problems of the premature infant. Ment Retard Dev Disabil Res Rev 2002; 8: 249–257.

[2] Wen SW, Smith G, Yang Q, Walker M. Epidemiology of preterm birth and neonatal outcome. Semin Fetal Neonatal Med 2004; 9: 429–435.

[3] B Larroque, G Bréart, M Kaminski, M Dehan, M André, A Burguet, H Grandjean, B Ledésert, C Lévêque, F Maillard, J Matis, J C Rozé, P Truffert, Survival of very preterm infants: Epipage, a population based cohort study, Arch Dis Child Fetal Neonatal Ed 2004;89:F139–F144.

[4] Jones, H.P., Karuri, S., Cronin, C.M., Arne Ohlsson, Abraham Peliowski, Anne Synnes, Shoo K Lee. Actuarial survival of a large Canadian cohort of preterm infants. BMC Pediatr 5, 40 (2005).

[5] Institute of Medicine. Preterm Birth: Causes, Consequences, and Prevention. Brief report. Washington DC, Institute of Medicine; 2006.

[6] Keshtkaran A, Keshtkaran V. Factors affecting neonatal death in Fars Province, Southern Iran, 2004. Midd East J Family Med 2007; 5: 42–45.

[7] Golestan M, Fallah R, Akhavan Karbasi S. Neonatal mortality of low birth weight infants inYazd, Iran. Iran J Reprod Med 2008; 6: 205–208.

[8] Khashu M, Narayanan M, Bhargava S, Osiovich H. Perinatal outcomes associated with preterm birth at 33 to 36 weeks’ gestation: a population-based cohort study. Pediatrics 2009; 123: 109–113

[9] Pierre-Yves Ancel, François Goffinet, Survival and Morbidity of Preterm Children Born at 22 Through 34 Weeks’ Gestation in France in 2011 Results of the EPIPAGE-2 Cohort Study, JAMA Pediatr. 2015;169(3):230–238.

[10] Souza, R.T., Costa, M.L., Mayrink, J, Francisco E. Feitosa, Edilberto A. Rocha Filho, Débora F. Leite, Janete Vettorazzi, Iracema M. Calderon, Maria H. Sousa, Renato Passini Jr, Philip N. Baker, Louise Kenny, Jose G. Cecatti, Perinatal outcomes from preterm and early term births in a multicenter cohort of low risk nulliparous women. Sci Rep 10, 8508 (2020).

[11] Ruthdel Río, MartaThió, Mattia Bosio, Josep Figueras, Martín Iriondo, Prediction of mortality in premature neonates. An updated systematic reviewPredicción de mortalidad en recién nacidos prematuros. Revisión sistemática actualizada, Anales de Pediatría (English Edition), Volume 93, Issue 1, July 2020, Pages 24–33

[12] Mekasha A, Tazu Z, Muhe L, Mahlet Abayneh, Goitom Gebreyesus, Abayneh Girma, Melkamu Berhane, Elizabeth M. McClure, Robert L. Goldenberg, Assaye K. Nigussie, Factors Associated with the Death of Preterm Babies Admitted to Neonatal Intensive Care Units in Ethiopia: A Prospective, Cross-sectional, and Observational Study. Global Pediatric Health. 2020, Volume 7: 1–9.

[13] Zhu Z, Yuan L, Wang J, Qiuping Li, Chuanzhong Yang, Xirong Gao, Shangqin Chen, Shuping Han, Jiangqin Liu, Hui Wu, Shaojie Yue, Jingyun Shi, Rui Cheng, Xiuyong Cheng, Tongyan Han, Hong Jiang, Lei Bao, Chao Chen. Mortality and Morbidity of Infants Born Extremely Preterm at Tertiary Medical Centers in China From 2010 to 2019. JAMA Netw Open. 2021;4(5):e219382.

[14] Feng, J., Lee, J., Vesoulis, Z.A. Fuhai Li. Predicting mortality risk for preterm infants using deep learning models with time-series vital sign data. *npj Digit*. Med. 4, 108 (2021).

[15] Li S, Gao J, Liu J, Hu J, Chen X, He J, Tang Y, Liu X and Cao Y (2021) Perinatal Outcomes and Risk Factors for Preterm Birth in Twin Pregnancies in a Chinese Population: A Multi-center Retrospective Study. Front. Med. 8:657862.

[16] Birhanu D, Gebremichael B, Tesfaye T, Tadesse M, Belege F, Godie Y, Wodaje M, Tamiru E. Survival status and predictors of mortality among preterm neonates admitted to neonatal intensive care unit of Addis Ababa public hospitals, Ethiopia, 2021. A prospective cohort study. BMC Pediatr. 2022 Mar 23;22(1):153.

[17] F. Gary Cunningham, Kenneth Leveno, Jodi Dashe, Barbara Hoffman, Catherine Spong, Brian Casey, Williams obstetrics. McGraw Hill / Medical; 26th edition (April 7, 2022)

[18] www.javatpoint.com

[19] Alka Rani, Nirmal Kumar, Jitendra Kumar, Jitendra Kumar, Nishant K. Sinha, Chapter 6 - Machine learning for soil moisture assessment, Deep Learning for Sustainable Agriculture, Cognitive Data Science in Sustainable Computing, 2022, Pages 143–168.

[20] www.freecodecamp.org

[21] www.geeksforgeeks.org

[22] www.towardsdatascience.com

[23] www.towardsdatascience.com

[24] www.medium.com

